# No Evidence of tACS-Primed Enhancement of Alpha Neurofeedback

**DOI:** 10.64898/2026.09.29.755297

**Authors:** Christopher Turner, Louis James, Mina Bakhtiari, Gemma Learmonth, Alessio Fracasso, Aleksandra Vuckovic

**Author notes:** **Corresponding authors:** Christopher Turner.

## Abstract

Neurofeedback shows promise as a clinical intervention, but its effectiveness is hindered by variability of response across individuals. One factor influencing neurofeedback success may be endogenous brain states, particularly the baseline power within target frequency bands. Transcranial alternating current stimulation (tACS) provides a potential mechanism to modulate these baseline frequency bands. This study explored whether tACS could effectively enhance participants’ ability to gain neurofeedback control by increasing lateralised posterior alpha activity prior to engagement in a covert spatial attention-based neurofeedback task. Participants completed three counterbalanced sessions: 1) alpha tACS and active neurofeedback, 2) alpha tACS and sham neurofeedback, 3) gamma (40Hz) tACS and active neurofeedback. Neurofeedback performance was assessed via lateralized posterior alpha power shifts and Posner task reaction times. We identified no difference in reaction times or lateralised alpha power shifts across conditions. While participants reliably demonstrated lateralised posterior alpha patterns, consistent with the covert spatial attention neurofeedback protocol, no advantage of active neurofeedback over sham was observed. The lack of significant effects underscores the need for further rigorous tACS studies to identify the parameters essential for successful neural entrainment, and to evaluate its potential in enhancing neurofeedback efficacy.

## 1 Introduction

Neurofeedback is a technique that provides users with real-time feedback of their ongoing brain rhythms, enabling them to actively control specific frequencies within the brain and bring about changes in cognitive and motor functions associated with these signals. Electroencephalography (EEG) is commonly employed for neurofeedback due to its low-cost, portability, and ease of use, requiring minimal setup and training. The high temporal resolution and extensive body of research linking oscillatory EEG patterns to cognitive processes has led to an increase in EEG-based neurofeedback research over recent years. Neurofeedback has been investigated as a potential intervention for a broad range of applications, including enhancing specific brain functions or behaviours in healthy adults (Enriquez-Geppert et al., 2013; Matsuzaki et al., 2023; Rydzik et al., 2023; Viviani & Vallesi, 2021), and addressing cognitive deficits and decline in clinical populations (Hong & Park, 2022; Marzbani et al., 2016; Patil et al., 2023; Vilou et al., 2023). However, the adoption of neurofeedback as a clinical intervention faces challenges due to inconsistent findings regarding its mechanisms and efficacy. A key factor contributing to these inconsistencies is the variability in participants’ ability to achieve volitional control over their EEG signals, with so-called ‘non-responder’ rates reported between 16% and 57% (Alkoby et al., 2018; E. Weber et al., 2011; L. A. Weber et al., 2020). This variability complicates the interpretation of weak or null findings, making it challenging to distinguish between the actual effectiveness of neurofeedback in modulating brain rhythms and inducing behavioural changes, and the possibility that participants are ineffectively engaging with the neurofeedback process itself. Furthermore, this uncertainty makes it difficult to translate findings to an intervention for broader populations, as it is difficult to determine those individuals who may be most likely to benefit.

The issue of non-responders in neurofeedback, and other interventions such as non-invasive brain stimulation, is an area of active investigation. One potential contributing factor is the brain state, or ongoing neural activity, prior to or during the application of the intervention. State-dependent effects have been identified as a contributing factor for the success of various types of brain stimulation (Benwell et al., 2015; Bradley et al., 2022; Learmonth et al., 2015; Schaworonkow et al., 2019). In the context of neurofeedback, pre-neurofeedback resting-state EEG, such as broadband eyes closed power (Nan et al., 2018), eyes closed alpha amplitude (Chikhi et al., 2023; Su et al., 2021; Wan et al., 2014), and resting state sensorimotor rhythm (Reichert et al., 2015), may serve as predictors of neurofeedback success. Specifically, these studies suggest that the higher resting-state power in the target neurofeedback frequency band predicts neurofeedback success. For example, Chikhi et al. (2023) report that participants who were more successful in regulating their EEG signals through neurofeedback had higher baseline power in the target (central-parietal alpha) band. However, they emphasize that this relationship is correlational and suggest that interventions capable of directly modulating brain rhythms, such as transcranial alternating current stimulation (tACS), could help clarify the causal link between resting-state power and neurofeedback success.

Transcranial Alternating Current Stimulation is a non-invasive electrical brain stimulation technique that applies alternating current to modulate neural activity by ‘entraining’ specific frequencies in targeted brain regions (De Koninck et al., 2023; Elyamany et al., 2021; Orendáčová & Kvašňák, 2021; Reato et al., 2013; Tavakoli & Yun, 2017). Given its ability to influence oscillatory activity, tACS may serve to optimise brain state for neurofeedback control by pre-conditioning neural rhythms to align with the target frequency. This approach remains largely unexplored, with only one known study integrating tACS and neurofeedback in this manner. Naros & Gharabaghi (2017) reported that intermittent beta-tACS enhanced classification accuracy in a task requiring self-regulation of neural rhythms to control an external device but with no lasting aftereffects of tACS. However, tACS-induced after-effects in the alpha frequency band are more commonly reported (Orendáčová & Kvašňák, 2021), and given that alpha based neurofeedback is a commonly used feedback protocol (Marzbani et al., 2016), the effect of tACS in priming alpha oscillations for neurofeedback warrants investigation.

In this study we aimed to investigate the potential priming effects of tACS stimulation on the ability of participants to perform a covert visual-spatial attention (CVSA) based EEG neurofeedback task. Past studies have consistently demonstrated the lateralisation of occipito-parietal alpha activity during CVSA tasks (Kornrumpf et al., 2017; Peylo et al., 2021; Thut et al., 2006; Worden et al., 2000). Specifically, these studies report a decrease in alpha power in the hemisphere contralateral to the attended location relative to the ipsilateral hemisphere, which is accompanied by faster reaction times to targets appearing in the attended hemifield. While this relationship between posterior alpha lateralisation and covert spatial attention (measured through reaction times to lateralised stimuli) is correlational, more recent studies using transcranial alternating current stimulation (tACS) to modulate posterior alpha activity have observed enhanced reaction times to cued, lateralised stimuli following stimulation (Kasten et al., 2016, 2020; Kemmerer et al., 2022; Middag-van Spanje et al., 2024; Radecke et al., 2023; Schuhmann et al., 2019). These findings suggest a causal link between posterior alpha oscillations and covert spatial attention. Building on this, we aimed to stimulate the occipito-parietal cortex at the alpha frequency to entrain occipito-parietal alpha activity, effectively ‘jump-starting’ participants’ ability to regulate these rhythms during neurofeedback. We hypothesised that entrainment via posterior alpha tACS would lead to a favourable brain state for subsequent CVSA-based neurofeedback training. Specifically, that active neurofeedback combined with tailored individual alpha frequency (IAF) tACS stimulation would improve the ability to control occipital-parietal alpha rhythms during subsequent active neurofeedback, relative to a sham neurofeedback condition.

Further, we hypothesise that this neurofeedback-driven lateralised alpha power increase will correspond to an improvement in reaction times during a subsequent Posner task measuring covert spatial attention.

## 2 Methods

### 2.1 Participants

Thirty adults (24 female, 29 right-handed) with ages ranging from 20 to 62 years (M = 27.26, SD = 9.13) were tested. All participants had normal or corrected to normal vision and completed a tACS safety screening form, which excluded people who were pregnant, had any metal fragments in their head, or were at risk of seizure. The study protocol was pre-registered on the Open Science Framework (https://osf.io/t8p5j/).

A sample size of 26 was calculated using G*Power (Faul et al., 2007) based on the effect size observed in a similar neurofeedback study (Schneider et al., 2020), with a view of performing a repeated measures ANOVA to compare the change in reaction time across three stimulation conditions (partial eta squared = 0.1, alpha = 0.05, power = 0.95). Four participants withdrew after their first session, one due to an inability to record eye tracking data through their glasses, and three due to scheduling conflicts; data from these participants were not included in the study. Of the remaining participants, one session was excluded for two participants (one active neurofeedback and gamma tACS, and one active neurofeedback and alpha tACS) due to missing reaction time and EEG data, resulting in 24 full datasets and two partial datasets in the final analysis (Mean age = 27.8, SD = 9.7). This study was approved by the University of Glasgow College of Medical, Veterinary & Life Sciences ethics committee and was performed in accordance with the Declaration of Helsinki, with all participants providing written consent prior to participation.

### 2.2 Experimental design

We adopted a single blind within-subjects design, where the experimenter was aware of the tACS/neurofeedback condition that was being administered during each session, but the participants were not informed. Each participant completed three sessions, with each on a separate day, and a minimum of 40 hours between sessions. During each session, the participants received one of the following protocols in a pseudo-randomised, counter-balanced order: a) alpha tACS and active neurofeedback; b) alpha tACS and sham neurofeedback; c) gamma (40Hz) tACS and active neurofeedback. Participants were not informed about the specific details of each session nor that the sessions would differ from one another. Fig. 2a describes the structure of each session which was identical regardless of the protocol the participant received. For each session, participants were seated in an electrostatically shielded room with the lights switched off and door closed during the tasks. Each task was displayed on a Samsung Syncmaster 1100MB monitor and fixation was maintained using a chin rest 60cm from the screen.

### 2.3 Transcranial Alternating Current Stimulation

During each session, either individual alpha frequency (IAF) or gamma frequency (40Hz) tACS stimulation was delivered with a NeuroConn DC Stimulator Plus device (neuroCare Group). Electrodes were attached to the Cz (vertex) and O1 (left primary occipital cortex) according to the EEG 10-10 system using Ten20 electrode paste. Electrode positions were identified to ensure an adequate electric field in the left occipital cortex while also ensuring neurofeedback could also be conducted with nearby electrodes (Fig. 1). We regularly monitored the electrodes to ensure that they did not move during the tasks. For alpha stimulation, each participant’s individual alpha frequency (*M* = 10.08Hz, *SD* = 1.07) was calculated from each session’s initial eyes-closed resting state EEG data using the *savgol_iaf* function from the Philistine python package (Alday et al., 2023), which is based on Corcoran et al. (2018). The IAF was calculated at the beginning of each session to account for variations in alpha frequency within individuals. Therefore, the specific frequency delivered could vary between sessions for each participant (*Mean SD =* 0.27Hz).

**Fig. 1.**
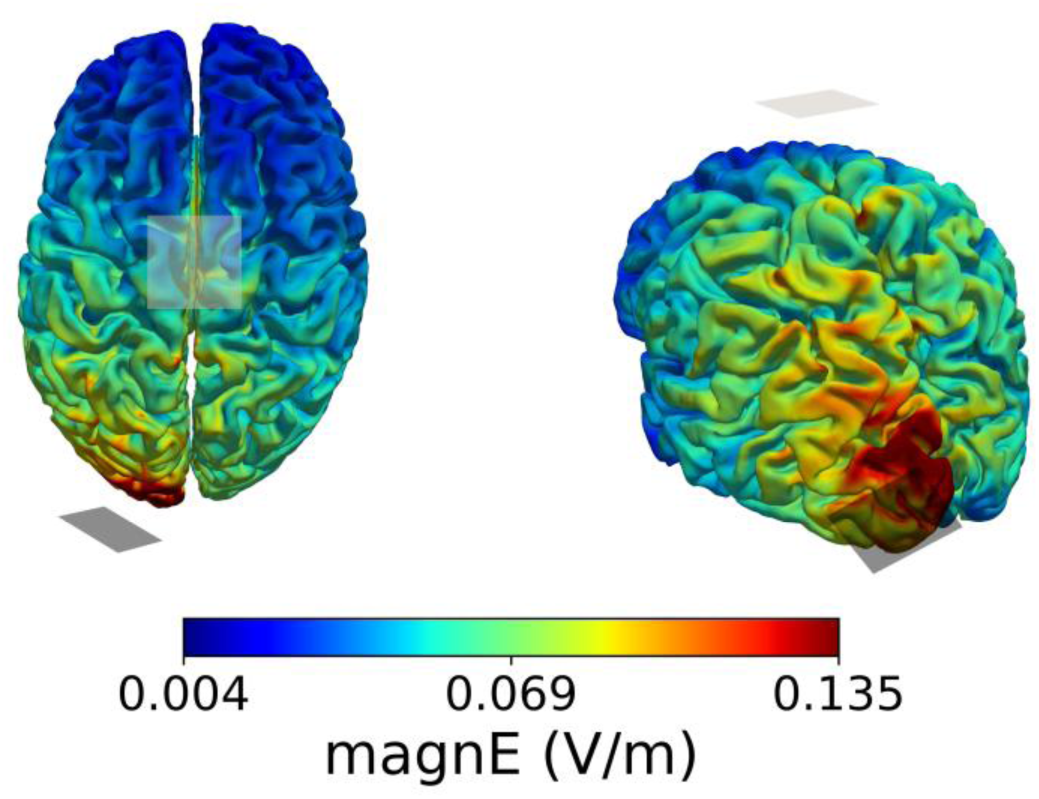
Simulated electric field distribution for the Cz–O1 tACS montage. Electric field magnitude (magnE, V/m) simulated using SimNIBS 4.6. Stimulation was delivered at 0.5 mA peak amplitude (1 mA peak-to-peak) with the anode at Cz and cathode at O1. Left: superior view. Right: posterior-inferior isometric view highlighting the occipital field distribution.

Stimulation was delivered after the first Posner task in a 15-minute continuous block with a one second ramp-up and ramp-down phase at the beginning and end, respectively. Using the method described in Vossen et al. (2015), the amplitude of the stimulation was calibrated to each participant’s sensitivity threshold. This was achieved by applying stimulation for 10 second periods at magnitudes starting at 1mA peak-peak and moving up in 0.25mA intervals until the participant; a) reported phosphenes, b) reported discomfort, or c) the maximum amplitude of 2mA peak-peak was reached. If phosphenes occurred or the stimulation created discomfort, the amplitude was reduced in 0.1mA steps until an acceptable level was achieved. This procedure was conducted once at the beginning of the first session with the calculated threshold used for subsequent sessions. The average current used across participants was 1.715mA (*SD* = 0.301). Each 10s calibration block was delivered with a 1 second fade in and fade out period to mimic the main stimulation block. EEG was not simultaneously recorded during the tACS delivery period.

### 2.4 Neurofeedback

The EEG neurofeedback task was implemented in the NFBLab software package (Smetanin et al., 2018). **Error! Reference source not found.**Participants completed a total of 12 trials, divided into four equal blocks, with each block separated by a participant-defined break. We instructed participants to maintain a fixed gaze on a black central fixation dot and to use covert visuospatial attention to modulate the colour of a left-lateralised target. The concept of covert attention was explained to participants prior to the experiment by telling them that they must keep their eyes fixed centrally while paying attention to something happening in their periphery. Participants were then given a chance to practice this at the beginning of each session with a shortened version of the neurofeedback task. Fig. 2c outlines the time course of each trial of the full neurofeedback task. Each trial began with a five second rest period indicated by a white central fixation dot displayed on a grey background. During this time the participant was able to rest their eyes and prepare for the training period. Following this, the central fixation dot changed colour to black, signalling the participant to focus on the centre of the screen. After 1s, two circular targets (3.5° diameter, vertically offset by -1° and horizontally offset by +/-5° visual angle) appeared on the screen indicating the start of the neurofeedback period. These targets appeared in identical positions as per the Posner task. The participant then had a two-minute window to control the colour of the left target with the corresponding right target kept black for the duration of the task. We instructed participants to alter the colour of the left target by aiming to maximise the intensity and duration of blue and minimise red. They were told that they could achieve this by shifting their covert attention to the left target, however, if they discovered their own strategies to achieve this, these could be used instead. The left target turned blue when the participant’s feedback signal was above an individualised threshold (see thresholding section below), and red when below. The intensity of the colour (blue or red) linearly increased from white (when at the threshold) to completely blue at the maximum, and red at the minimum values.

**Fig. 2.**
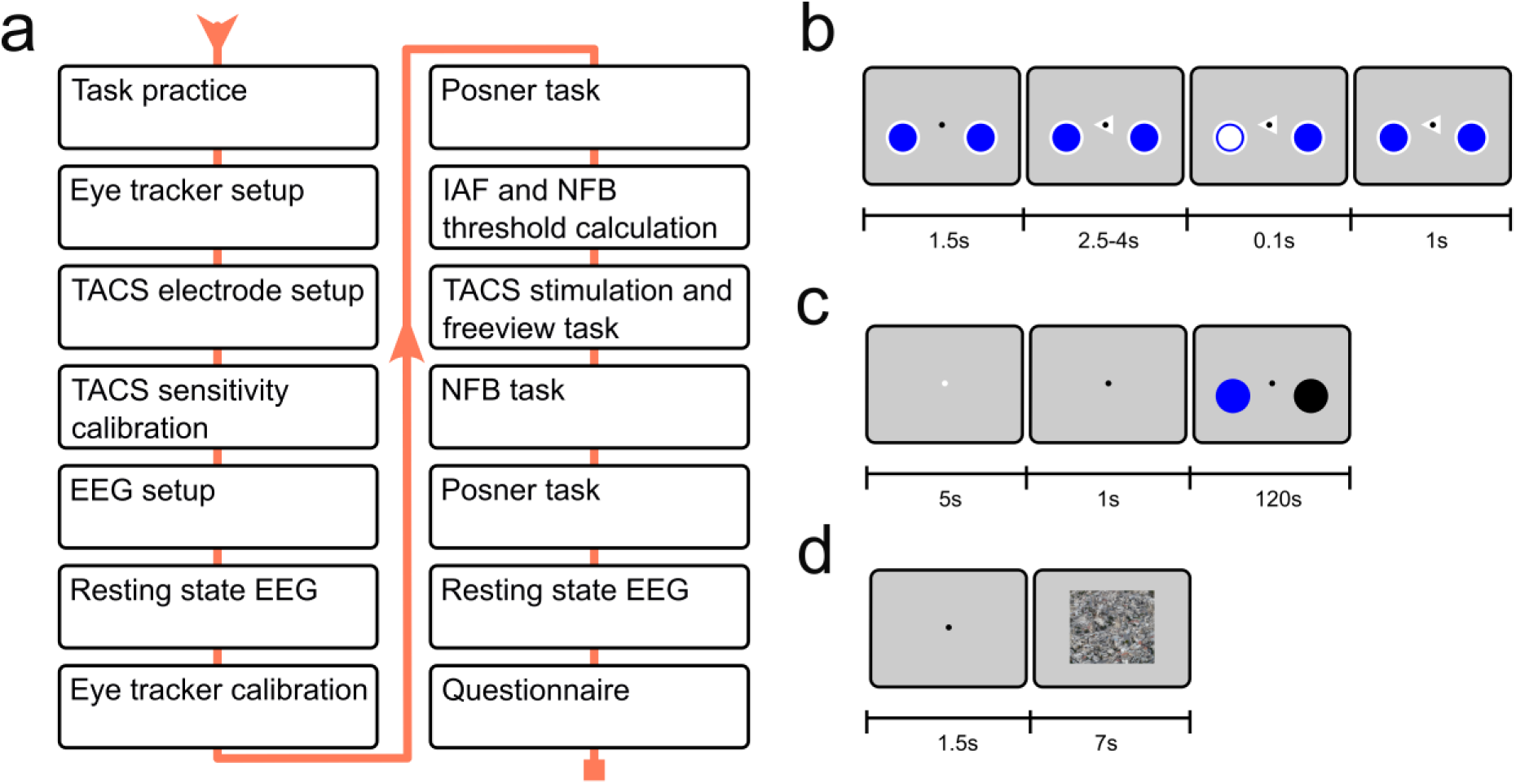
Session structure and task trial structure. **a)** Structure for each individual session starting from the top left and ending at the bottom right. Note, tACS=transcranial Alternating Current Stimulation, IAF=individual alpha frequency, NFB=neurofeedback. **b)** Posner task trial structure. **c)** Neurofeedback task trial structure. The left target dynamically changed from blue to red depending on the alpha lateralisation of each participant (for active neurofeedback), or from a pre-recorded session (sham neurofeedback). The right target remained black for the duration of the experiment. **d)** Freeview task trial structure.

In the active feedback session, the colour of the left target was controlled by the alpha asymmetry index (AAI) signal from the P5 and P6 electrodes. The neurofeedback AAI signal is described as *(α_P5_* -*α_P6_)/ (α_P5_* + *α_P6_)* where *α_P5_* is the alpha power from the P5 electrode (ipsilateral to shifting covert attention to the left) and *α_P6_* is the alpha power from the P6 electrode (contralateral to shifting covert attention to the left). Consequently, a positive AAI value represents higher left (ipsilateral) alpha power relative to right (contralateral) alpha power. During sham neurofeedback, the colours were controlled in the same manner but a recording from a different participant was used to calculate the online AAI (i.e., the participant had no direct control over the colours).

Alpha power at P5 and P6 was calculated online at each participant’s individual alpha frequency (IAF) using the NFBLab software with *FFT envelope detection* and a 1000-sample window. A 100-sample moving-average filter was applied to the online AAI signal to reduce jitter and make the colour transitions appear smoother to the participant.

#### Thresholding

The neurofeedback threshold was calibrated from EEG data collected during the first Posner task at the beginning of each session. All AAI power values from left and right cued trials were fit to a normal distribution with the mean used to set the threshold and the maximum and minimum values set to +/- 2 standard deviations, respectively. A constant offset of 20% of the range (maximum – minimum values) was applied to the calculated mean to control for natural variation in the AAI value and induce learning during the neurofeedback task, i.e. if the participant was at rest, the left target would predominately remain red.

### 2.5 Posner task

We measured reaction times before and after the tACS and neurofeedback interventions via a Posner task, implemented in Python v3.8 and PsychoPy 2022.2.4 (Peirce et al., 2019). Each Posner task consisted of 100 trials divided into four equal blocks of 25 trials, with each block separated by a participant-defined break. Individual trials began with a 1.5s fixation period where a black central fixation dot was displayed on a grey background (Fig. 2b). Two blue circles (3.5° diameter, vertically offset by -1° and horizontally offset by +/-5° visual angle) to the left and right of the central fixation point were always present on the screen and represented the two possible locations of the upcoming target stimulus.

Following the fixation period, a directional cue was displayed in the form of a white triangle, centrally positioned behind the black fixation dot. The cue could either point to the left, right, or both directions with an equal probability. After a pseudo-random delay between 2.5 and 4 seconds, the target stimulus (a white circle, 1° visual angle diameter) appeared for 100ms in the centre of either the cued (valid) or un-cued (invalid) targets with a validity of 70%. A response period of 1s followed the stimulus where the participant was required to press either the left arrow key (for left stimuli) or right arrow keys (for right stimuli) on a keyboard centred in front of them. We instructed participants to maintain fixation on the central fixation dot throughout the trial and to respond to the appearance of the stimuli with their dominant hand as fast and accurately as possible.

### 2.6 Free-viewing Task

During tACS stimulation, participants performed a free-viewing task based on Chiffi et al. (2021). This was a filler task to ensure that participants were engaged in a visual task during the tACS period, but no responses were required. The task was started manually by the experimenter at the same time as the tACS onset and consisted of four blocks of 24 trials separated by an eye-tracker drift correction. Each trial consisted of a black central fixation dot followed by an image displayed for 7 seconds (Fig. 2d). Participants were instructed to maintain their gaze during the fixation period and freely explore the image when it appeared. The images were centrally displayed with a height and width corresponding to 50% of the screen dimensions. These images were selected from a set of 120 from Chiffi et al. (2021), originally sourced from the free image database Pixabay (pixabay.com) from the subcategories “nature” and “architecture”, and used with the author’s permission. Each image was shown either in its original orientation or mirrored horizontally, with participants viewing an image or its mirror only once per session. An algorithm was used to ensure balanced saliency between the left and right of all images (Chiffi et al., 2021). The results of this task were not analysed for the present study.

### 2.7 EEG Acquisition and Pre-processing

A brainAmp MR (Brain Products GmbH) with a 64 channel brainCap was used to record the EEG data from passive electrodes according to the international 10-10 system.

The EEG cap was placed over the attached tACS electrodes, taking care not to alter their positions. Electrodes O1 and Cz were not recorded as these were over the sites of the tACS electrodes. Electrodes sites were cleaned and prepared with Abralyt (Easycap GmbH) abrasive gel. EEG recordings were acquired at a sampling rate of 1000Hz with the online reference electrode at FCz and the ground at AFz. One electrode was placed on the right outer canthus to assist in removing large eye artefacts from the EEG data. We used BrainVision Recorder software to visually check the signal quality during setups and aimed to keep impedances below 10kΩ. After setup, all participants completed an initial eyes-open and eyes-closed resting state EEG recording (two minutes each) where they were asked to sit at rest. This resting state EEG was also recorded after the last neurofeedback task and was repeated in every session. BrainVision Recorder was used to record the EEG during the Posner tasks, and NFBLab (Smetanin et al., 2018) was used to record EEG data during resting state and neurofeedback tasks. This distinction reflected the practical setup of the experiment, as NFBLab was required for the neurofeedback sessions but was not used during the Posner tasks.

Offline, pre-processed EEG data was used for all analyses. Preprocessing was conducted with MNE Python software (Gramfort et al., 2013). First, the tACS electrodes (O1, Cz) were excluded, then the raw data was filtered between 1 and 40Hz separately for every task in each session. Each of the filtered EEG data files were then concatenated per session and used as input for Independent Component Analysis (ICA), using the fastICA algorithm (Hyvarinen, 1999), where components representing eye blinks, lateral eye movements, and muscle artefacts were visually assessed and removed from the data. Finally, epochs with an instantaneous voltage exceeding 100uV were removed. For neurofeedback, epochs were obtained by splitting the signal into 1 second, non-overlapping, segments. For the Posner tasks, epochs were defined as whole trials (i.e., 6.5s). The AAI signal used during online feedback was re-calculated from offline data using the same functions from the NFBLab software package. This was calculated for each session. No re-referencing of the EEG data was conducted to obtain the signal that participants used during feedback.

### 2.8 Eye tracking

We recorded binocular eye position using an SR Research Eyelink 1000 eye-tracker with a sampling rate of 500Hz. An initial setup and 5-point calibration routine was performed before attaching the stimulation or EEG electrodes to ensure suitability of eye tracking for each participant. The eye tracker was calibrated before starting each task, and before each neurofeedback block. We performed drift corrections between blocks in the Posner and tACS tasks. Eye-tracking was used to control participant gaze and ensure central fixation during the tasks. Results of eye traces analyses will be presented elsewhere.

#### Questionnaires

After each session, participants completed questionnaires regarding their experiences during the session (available on OSF, https://osf.io/t8p5j/). Specifically, we measured participants’ motivation prior to the session, their expectations for achieving neurofeedback control, any strategies they used, their subjective sense of success with the neurofeedback tasks after the session, and an open section for further comments and feedback. After the final session, participants were asked if they could identify which neurofeedback session was sham and were then unblinded to the experimental conditions. We also asked participants to rate the strength and duration of any sensations during tACS stimulation (headache, tingling, itching, burning, or pain) they may have experienced at the end of each session.

### 2.9 Statistical Analysis

All statistical tests aside from those used to analyse the questionnaire data were conducted using a linear mixed effects model in *R 4.3.1* (R Core Team, 2023) with the *lmer* function from the *lme4* package (Bates et al., 2015) with *p-*values obtained using the *anova* function in the *lmerTest* package (Kuznetsova et al., 2017). Bonferroni corrected post-hoc comparisons were calculated with the *emmeans* package (Lenth, 2023). Bayes Factor analysis was conducted with the *BayesFactor* package (Morey & Rouder, 2022) and was used to compare models and determine the level of evidence for any significant effects. Normality of residuals was visually determined with a q-q plot (*ggqqplot* from the *ggpubr* package (Kassambara, 2023a) and confirmed with Shapiro-wilk tests using the *shapiro_test* from the *rstatix* package (Kassambara, 2023b). Each of these models was fitted twice: once with session coded by intervention protocol (active neurofeedback with alpha-tACS, active neurofeedback with gamma-tACS, and sham neurofeedback with alpha-tACS), and once by session order (first, second, and third session). Both codings have three levels.

#### 2.9.1 Reaction Times

We first analysed valid and invalid Posner trials directly to uncover any validity effect (differences in median reaction times between valid and invalid trials). Participants’ response times are generally slower in invalid compared to valid trials (Posner, 1980), and so slower reaction times would be expected in invalid trials compared to valid in our task. Further, the validity effect was analysed both by *session protocol* (i.e. the different intervention combinations), and *session number* (i.e. the order in which sessions were completed) to investigate any learning effects over the three sessions. A linear mixed effects model was used with task (two levels: pre-and post-intervention), session (three levels), cue validity (two levels: valid and invalid), and their interactions modelled as fixed effects with random intercepts for each participant.

The pre-registered analysis plan defines the primary outcome measure as change in lateralised reaction times for valid cues(*ΔLRT_v_*) between the pre- and post-intervention tasks. However, to increase statistical power, we have used the (*LRT_v_*) for all tasks and included task (pre- and post-intervention) as a factor in the model. *LRT_v_* is defined as *RT*_*left_v_* − *RT*_*rig*ℎ*t_v_* where *RT*_*left_v_* and *RT*_*rig*ℎ*t_v_* are the median reaction times for left and right valid trials respectively for each participant. Note, to preserve units, we have not used the normalised version of *LRT_v_* as described in the pre-registration. The *LRT_v_* for each participant was calculated for the pre-and post-intervention tasks for each session. A linear mixed effects model was used to test for effects of task (two levels: pre- and post-intervention), session (three levels), and their interaction, modelled as fixed effects with random intercepts for each participant.

#### 2.9.2 Alpha Asymmetry Index

The EEG outcome measure was the median AAI during each trial. The mean AAI for the neurofeedback task was calculated for each two-minute window of active neurofeedback. For the Posner task, this was calculated for the cued period (from 1.5s to 4s after trial start). Note that the analysis of Posner electrophysiological data was not outlined in the pre-registration, however we have included this as an exploratory analysis to investigate any potentially lasting effects of the interventions on alpha power. For the neurofeedback data, task (six levels: two eyes-open resting-state blocks and four neurofeedback blocks), session (three levels), and their interaction were modelled as fixed effects, with participant intercepts as random effects, in a linear mixed effects model. For the Posner data, task (two levels: pre-and post-intervention), session (three levels), cue direction (three levels: left, right, and centre), and their interactions were modelled as fixed effects, with participant intercepts as random effects.

For completeness, we show the topography of alpha power (8-12Hz) along with the time frequency plots for the P5 and P6 electrodes for the Posner and neurofeedback trials (Fig. 3). For the Posner task, the topography shows the normalised difference between left and right cues, i.e., 100*(left-right)/(left+right). For the neurofeedback task, since there are no left and right cued trials, the topography shows the change in alpha power above the mean of a baseline period (-250 to -50ms) before the beginning of the trial. The time frequency plots for the neurofeedback task show the change in log-transformed power above the baseline period (-250 to -50ms) before the beginning of the trial.

**Fig. 3.**
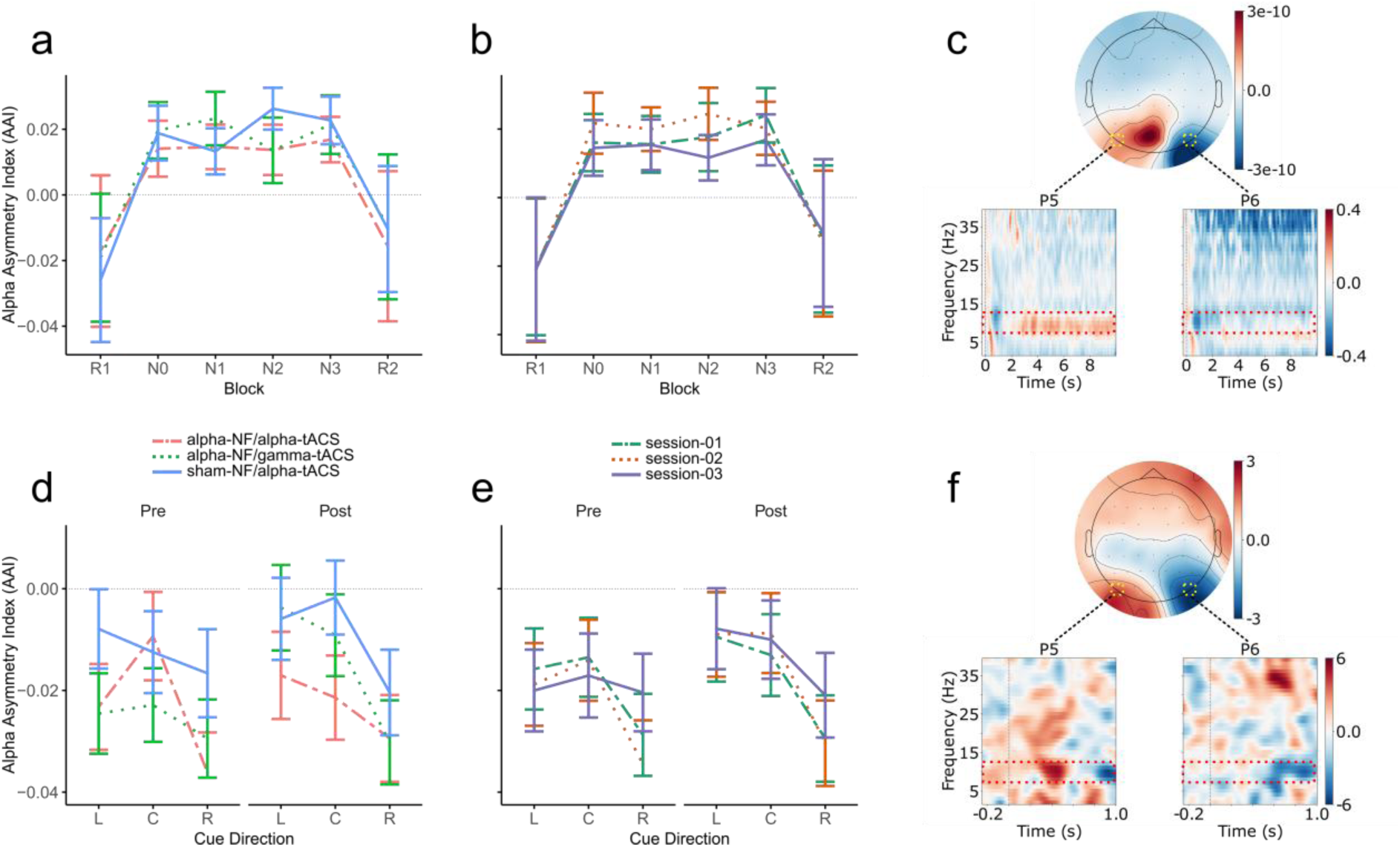
Alpha asymmetry results. **a)** Mean AAI averaged over participants for each session protocol, for each resting state (pre-session eyes-open (R1), and post-session eyes-open (R2)), and neurofeedback block (block0 (N0), block1 (N1), block2 (N2), block3 (N3)). **b)** Mean AAI averaged over participants for each session, for each resting state and neurofeedback block. **c)** Topographic plot of alpha power during neurofeedback trial (8-12Hz) baselined with the period of -250 to -50ms. Time-frequency plots for the P5 and P6 electrodes show change in log-transformed alpha power over the baseline period -250 to -50ms. The red dotted outline shows the alpha band and time 0s indicates the start of the neurofeedback trial. All error bars show 95% confidence intervals. **d)** Mean AAI averaged over participants for each session protocol: active-NF/alpha-tACS, active-NF/gamma-tACS, sham-NF/alpha-tACS; for each cue direction: Left (L), Centre (C), and Right (R); for the pre-and post-Posner tasks. **e)** Mean AAI averaged over participants for each session for each cue direction and Posner task. **f)** Topographic plot of normalised difference of left and right cue alpha power during the Posner trials (8-12Hz) with time-frequency plots for the P5 and P6 electrodes. The vertical dotted line shows the onset of the cue, and the red dotted outline shows the alpha band.

#### 2.9.3 Trial Exclusion Criteria

Eye-tracking was used to control participant gaze and ensure central fixation during the tasks. Specifically, we only included trials with valid eye tracking data in our analysis.

Valid eye tracking data was defined as timepoints where the gaze position was not lost (due to eyes closing, blinks, and other factors), and with a gaze position within the inner horizontal bounds of the left and right targets i.e., when participants were centrally fixating. Trials were excluded according to the following criteria: For the neurofeedback task, a whole trial was excluded if the number of valid eye tracking time points was less than 25% of the total trial. Four (0.4%) neurofeedback trials were removed leaving a total of 920 trials across all participants. For the Posner task, a trial was excluded if the number of included time points was less than 50% of the trial. Additionally, an entire Posner task was excluded from analysis if more than 50 trials (50%) were removed due to bad eye data. In total, one task was excluded across participants. Posner task trials were also excluded due to behavioural performance. Specifically, only trials where a single, correct response was given between 200ms and 1000ms were included in the analyses. A task was excluded if less than 75% of the responses were correct. Overall, 650 Posner trials (4%) were removed leaving a total of 14250 trials included in the analysis.

#### 2.9.4 Questionnaires

Questionnaire analysis was not included in the pre-registration protocol; however, we conducted an exploratory analysis of the questionnaire data, specifically, on measures of motivation and perceived success. All statistical tests were conducted in *R 4.3.1* (R Core Team, 2023). We conducted a correlation between these two measures with a Bayesian Spearman rank correlation using the *correlationBF* function from the *correlation* package (Makowski et al., 2025) to determine evidence for the null or alternative hypotheses. We further compared the participants’ feeling of subjective success after the sessions with a Kruskal-Wallis H test using the default *kruskal.test* function. Additionally, we compared the difference in prior motivation and subjective success for participants who correctly identified the sham session with those who did not, separately with a Kruskal-Wallis H test. To investigate if participants could identify the sham session more than chance, we conducted a two-sided binomial test on the participants’ correct sham guess (true/false) with ‘true’ having a 1/3 chance (as there was one sham neurofeedback session in three sessions) of being answered and ‘false’ a 2/3 chance. The binomial test was conducted using the *binom.test* function. A Bayes factor was further calculated with the *proportionBF* function from the *BayesFactor* package (Morey & Rouder, 2022) to determine if sufficient for the null and alternative hypotheses. We used a cumulative link mixed model (CLMM) to assess whether adverse sensation severity differed across stimulation sessions. Severity was rated on a 5-point ordinal scale, and the model included session protocol as a fixed effect, with a random intercept for each participant. The model was estimated using the Laplace approximation via the *clmm* function in the *ordinal* R package (Christensen, 2024). Finally, we descriptively report the strength and duration of tACS stimulation reported by participants.

## 3 Results

### 3.1 Neurofeedback

The negative AAI values during the eyes-open rest shown in Fig. 3a indicate that overall, participants had higher alpha power in the right hemisphere compared to the left during these blocks. However, when participants were actively performing the neurofeedback task, the positive AAI values indicate that relative alpha power shifted to the left hemisphere. Notably, this leftward shift is comparable in both active and sham conditions. A linear mixed effects model showed a significant main effect of task (F(5, 418.06) = 18.13, *p* < 0.001) with post hoc tests indicating significantly higher AAI (i.e., higher alpha power in the left hemisphere compared to right) in all neurofeedback blocks compared to resting-state (*p* < 0.0001). However, there were no differences within individual neurofeedback blocks, or within individual resting-state blocks. There was also no interaction with task and session protocol, indicating no differences between each of the session protocols. Q-Q plots and Shapiro-wilk tests indicated that the residuals were normally distributed (p > 0.05). A Bayes Factor analysis indicated evidence for a model with no interactions (BF = 4.82e-3 ± 1.86%), and evidence for a model including task (BF = 2.52e11 ± 1.92%).

Similar results are observed when looking at session order (Fig. 3b). There was a main effect of task (F(5, 418.06) = 18.04, *p* < 0.001) with post-hoc tests showing AAI in all neurofeedback blocks significantly left-shifted compared to those in the resting-state tasks (*p* < 0.0001). However, there were no differences within neurofeedback blocks or within resting-state blocks. Q-Q plots and Shapiro-wilk tests indicated that the residuals were normally distributed (*p* > 0.05). A Bayes Factor analysis indicated evidence for a model with no interactions (BF = 2.67e-3 ± 1.95%), and evidence for a model that includes task (BF = 1.39e11 ± 1.37%).

The topography plot in Fig. 3c shows posteriorly lateralised alpha power, with higher alpha power in the left hemisphere relative to the right (the neurofeedback target, and thus the focus of covert spatial attention, was on the left side during this task). The time-frequency plots for the neurofeedback electrodes P5 and P6 show a maintained, consistent increase in log-transformed alpha power over the baseline period (-250 to -50ms) in the alpha band (8-12Hz) for the left electrode (ipsilateral to directed covert attention) and a slight decrease in alpha power for the contralateral right electrode.

### 3.2 Posner Task

#### 3.2.1 Alpha Asymmetry

The AAI analysis (alpha values at electrodes P5 minus P6) for the Posner task revealed no interactions but a marginally significant main effect of session protocol (F(2, 410.64) = 3.09, *p* = 0.05). However, a post hoc pairwise comparison using estimated marginal means revealed no significant differences between any of the session protocols. There was also a main effect of cue direction (F(2, 410.01) = 6.59, *p* = 0.002) with post hoc estimated marginal means tests identifying significantly smaller AAI values only when comparing right cues to centre cues (*p* = 0.005) and right cues to left cues (*p* = 0.006).That is, when compared to centre and left cues, right cued trials elicited a reduction in alpha power in the left hemisphere compared to right; whereas centre and left cues induced similar alpha power in both hemispheres. Q-Q plots and Shapiro-wilk tests indicated that the residuals were normally distributed (*p* > 0.05). The Bayes Factor analysis of the full model compared to a model with no interactions suggests evidence in support of the model with no interactions (BF = 1.7e-05 ± 6.52%). However, when comparing the full model to a model with no cue direction, there is evidence for rejecting the model including cue direction (BF = 1.84e-4 ± 4.02%). When comparing to a model with no session protocol, there is evidence to reject the model including session protocol (BF = 7.99e-06 ± 3.37%). Fig. 3d shows the grand average AAI values for each session protocol recorded for each cued direction in the pre-and post-Posner task. When considering a learning effect (Fig. 3e), there are no interactions and only a main effect of cue direction (F(2, 410.01) = 6.37, *p* = 0.002). As with session protocol, post hoc tests show significantly more rightward (negative) AAI values for right cues when compared to centre cues (*p* = 0.006) and right cues compared to left cues (*p* = 0.007). Q-Q plots and Shapiro-wilk tests indicated that the residuals were normally distributed (*p* > 0.05). The Bayes Factor analysis shows evidence in support of a model with no interactions (BF = 1.96e-06 ± 7.29%), and evidence in support of a model without cue direction (BF = 2.03e-05 ± 3.43%).

The topographical plot showing normalised difference in alpha power between left and right cues shows the typical posterior lateralisation with lower posterior alpha power contralateral to the cue direction (Fig. 3f).

#### 3.2.2 Reaction Times

When comparing reaction times between the session protocols, participants had reliably faster reaction times for valid cues compared to invalid (Fig. 4a). A linear mixed effects model showed a significant main effect of cue validity (F(1, 267.00) = 237.75, *p* < 0.001) with *emmeans* post-hoc tests showing faster reaction times to valid cues (*p* < 0.0001). Additionally, we observed a main effect of session protocol (F(2, 267.46) = 3.30, *p* = 0.04) with Bonferroni corrected post-hoc tests indicating generally faster reaction times for active-NF/gamma-tACS when compared to the sham-neurofeedback with alpha tACS (sham-NF/alpha-tACS) condition only (*p* = 0.047). There were no significant interactions. Q-Q plots and Shapiro-wilk tests indicated that the residuals were normally distributed (*p* > 0.05). A Bayes Factor analysis with the full model compared to a model with no interactions indicated evidence in support of the no interactions model (BF = 1.0e-4 ± 4.98%). When considering the full model compared to a model with no interactions and no cue validity, there is evidence of a main effect of cue validity (BF = 3.8e32 ± 5.94%). However, when considering the full model compared to a model omitting interactions and session protocol there is evidence of no effect of session protocol (BF = 8.5e-05 ± 2.89%).

**Fig. 4.**
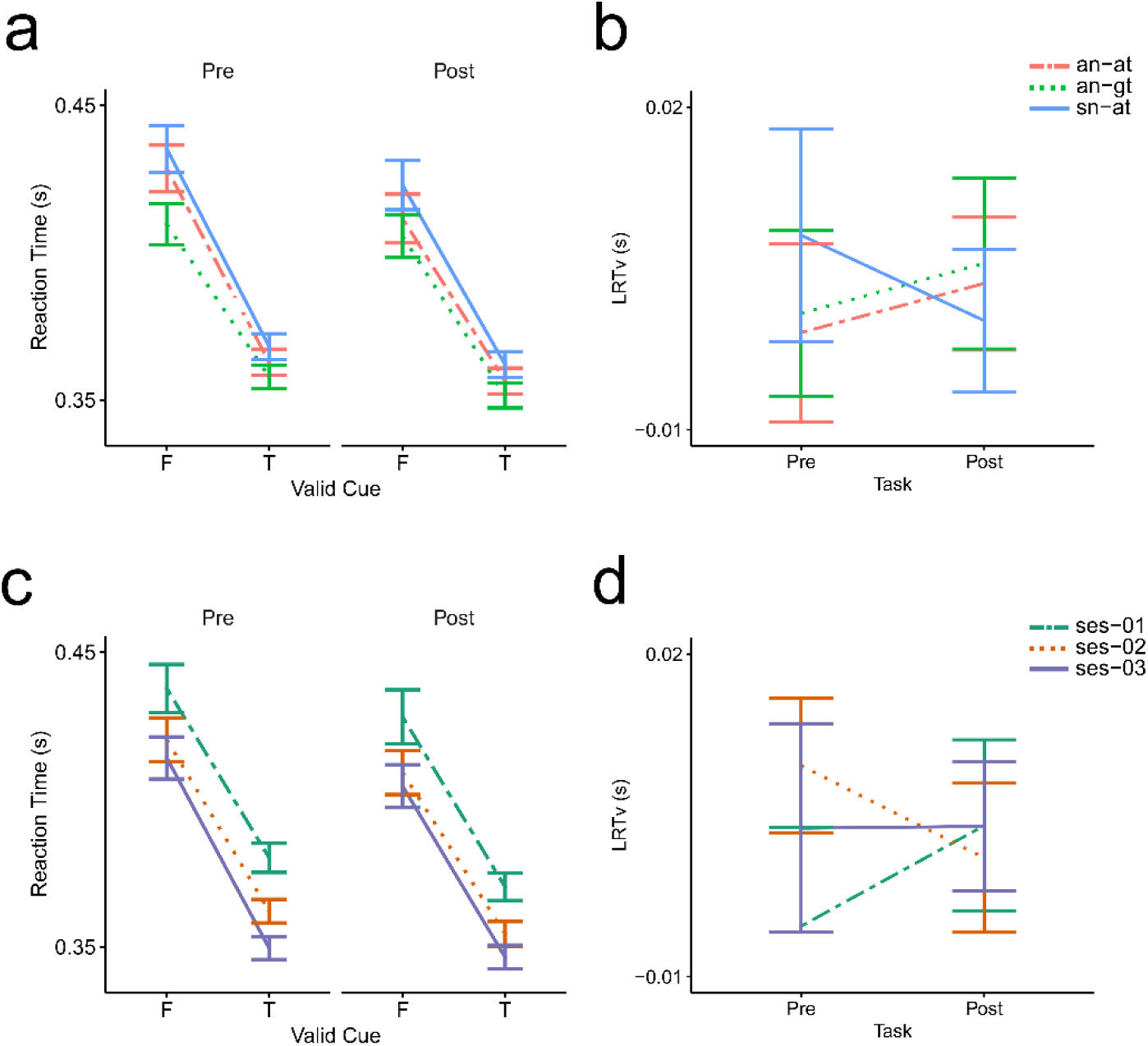
Reaction time results from the Posner task. **a**) Mean reaction times, averaged across participants for each session protocol: active-NF/alpha-tACS, active-NF/gamma-tACS, and sham-NF/alpha-tACS. Plots show faster reaction times for valid cues (T) compared to invalid (F) and similar results for the pre- and post-Posner tasks. **b)** Lateralised reaction times for valid cues (LRTv) averaged across participants for each session protocol. Plots show no change in lateralised reaction times between the pre- and post-Posner tasks. **c)** Mean reaction times, averaged across participants for each session, independent of the protocol: session 1 (ses-01), session 2 (ses-02), session 3 (ses-03). Plots show faster reaction times for valid cues (T) compared to invalid (F) and similar results between the pre- and post-Posner tasks. **d)** Lateralised reaction times for valid cues (LRTv) averaged across participants for each session. Plots show no change in lateralised reaction times between the pre- and post-Posner tasks. All error bars show 95% confidence intervals

When investigating a possible learning effect across the 3 sessions, participants similarly had generally faster reaction times for valid compared to invalid cues (Fig. 4c). There was a significant main effect of session number (F(2, 267.40) = 19.70, *p* < 0.001), and cue validity (F(1, 267.0) = 266.68, *p* < 0.001). Post hoc tests revealed faster reaction times for valid compared to invalid cues (*p* < 0.0001), and faster reaction times for both the second and third sessions when compared to the first (*p* < 0.0001) but no difference in reaction times between the second and third sessions. This indicates that participants quickly learned to improve their reaction times globally after the first session but did not improve further after the second. No significant interactions were identified. Q-Q plots and Shapiro-wilk tests indicated that the residuals were normally distributed (*p* > 0.05). The Bayes Factor analysis of the full model compared to a model with no interactions shows evidence in support of the model with no interactions (BF = 1.85e-4 ± 7.63%). Further when comparing the full model to a model omitting interactions and session number, there is evidence in support of including session number (BF = 291.04 ± 12.95%). Likewise, when comparing the full model to a model omitting interactions and cue validity, there is evidence in support of including cue validity in the model (BF = 7.76e35 ± 9.48%).

Overall, there was no difference in lateralised reaction times for valid targets between the pre-intervention and post-intervention tasks with no significant main effects or interactions. This can be also seen by the overlapping confidence intervals and the small magnitude of lateralised valid reaction times when comparing session protocol (Fig. 4b) and session number (Fig. 4d). Q-Q plots and Shapiro-wilk tests indicated that the residuals were normally distributed (*p* > 0.05). However, when comparing the full model to a model with no interactions, there is only anecdotal evidence of the absence of an effect for both session number (BF = 1.01 ± 3.7%) and session protocol (BF = 0.47 ± 2.8%).

### 3.3 Questionnaires

A Bayesian Spearman rank correlation between subjective success (M = 4.2, SD = 1.2) and prior motivation (M = 5.8, SD = 0.9) indicated only anecdotal evidence for the alternative hypothesis (rho = 0.22 BF = 2.35 ± 0%).

Kruskal-Wallis rank sum tests showed no significant differences between the subjective success of participants between session number or session protocol. Additionally, Kruskal-Wallis rank sum tests showed no significant difference of subjective success or prior motivation for people who correctly identified the sham neurofeedback session compared to those who did not.

Half the participants (13/26) correctly identified their sham neurofeedback session. A two-tailed binomial test indicated that this was not above chance (where chance is 1/3, *p* = 0.09). Furthermore, the Bayes factor of 1.45 ± 0.01% indicates only anecdotal evidence for the alternative hypothesis.

Finally, Table *1* indicates the number of sessions where participants reported tACS sensation strength and duration. A cumulative link mixed model (CLMM) showed no significant differences in sensation strengths or durations across the 3 protocols.

**Table 1.** Number of sessions with reported adverse tACS sensations.

| Strength | headache | tingling | itching | burning | pain |
| --- | --- | --- | --- | --- | --- |
| 1 (not at all) | 65 | 26 | 50 | 63 | 70 |
| 2 | 11 | 19 | 15 | 5 | 7 |
| 3 | 2 | 12 | 9 | 6 | 1 |
| 4 | 0 | 14 | 3 | 3 | 0 |
| 5 (very strongly) | 0 | 7 | 1 | 1 | 0 |
| Duration |  |  |  |  |  |
| No sensation | 62 | 27 | 51 | 63 | 68 |
| A few seconds | 6 | 17 | 8 | 8 | 5 |
| 1-2 mins | 1 | 12 | 5 | 3 | 2 |
| 3-5 mins | 4 | 1 | 1 | 4 | 1 |
| 6-10 mins | 1 | 9 | 4 | 0 | 1 |
| 11+ mins | 3 | 6 | 3 | 0 | 1 |

*Note.* Each value represents the number of sessions (maximum n = 78) where participants reported the sensation.

## 4 Discussion

In this study we investigated the effects of transcranial alternating current stimulation (tACS) at individual alpha frequencies on performance during a single session of covert spatial attention-based neurofeedback. Our findings indicate that IAF-tACS had no effect on neurofeedback performance as there were no differences in alpha asymmetry or Posner task reaction times between the alpha and gamma tACS protocols. Further, no differences in reaction times were observed between the pre- and post-stimulation Posner tasks. However, as expected, we observed a validity effect, with faster reaction times in response to valid Posner cues compared to invalid cues, regardless of session order, session protocol, or timing of the Posner task (pre- and post-intervention). Additionally, we observed a learning effect for reaction times across sessions, where participants had globally faster reaction times across all cue types following the first session, with performance plateauing in subsequent sessions. Finally, no evidence was found for a correlation between participants’ self-reported neurofeedback success and their motivation levels prior to the session. Participants were also unable to identify the sham neurofeedback session at a level greater than chance, indicating effective sham blinding.

### 4.1 Neurofeedback Does Not Impact Behavioural Results or Lateralised Alpha Power

Our primary aim was to ‘prime’ the endogenous cortical state using alpha tACS, prior to administering alpha neurofeedback, with the aim of improving the efficacy of the neurofeedback intervention. Our results showed that participants had increased leftward lateralisation during neurofeedback blocks compared to the eyes-open baseline, consistent with engagement of covert attention to the left side of space. Inspection of the time– frequency data (Fig. 3c) suggests that this shift is predominantly driven by increased alpha power in the left (ipsilateral) hemisphere, with a concurrent but smaller decrease in right (contralateral) alpha power. However, no differences were found between active and sham neurofeedback, pre- and post-resting states, or within individual neurofeedback blocks.

These findings could suggest several possibilities. Firstly, the lower alpha power in the right parietal cortex during the neurofeedback task indicates that participants were performing the left covert spatial attention task successfully (Bahramisharif et al., 2010; Samaha et al., 2016). Further, the posterior alpha lateralisation observed during the resting state periods show typical resting-state lateralisation patterns, with alpha power higher in the right parietal cortex relative to the left, consistent with previous findings that resting-state alpha activity reflects intrinsic hemispheric asymmetries in attentional and perceptual processing (Barry et al., 2007; Shulman et al., 2010; Spagna et al., 2020). However, despite this clear engagement with the covert attention task during neurofeedback, the neurofeedback intervention itself did not improve task performance any further. Schneider et al., (2020) similarly found no overall effects of neurofeedback in a covert spatial attention-based protocol, although they reported trends towards neurofeedback learning and significant improvements in performance for leftward trials. In a separate study, Tonin et al., (2013) reported significant above-chance classification accuracy in a comparable CVSA-based brain computer interface (BCI) paradigm, indicating successful control of the system via covert spatial attention. Conversely, we found no learning effects across blocks or sessions, suggesting that participants did not improve in their ability to modulate alpha lateralisation with practice, or alternatively, were already performing at a stable level from the outset. A key difference between our study and Schneider et al., (2020) and Tonin et al., (2013) is that we exclusively presented the neurofeedback target stimulus on the left side of space. Constantly training attention in a single direction may have contributed to a ceiling effect, similar to other neurofeedback studies (Biswas & Ray, 2019). However, in the absence of learning effects across blocks or sessions, it may also be possible that the neurofeedback paradigm employed here was simply ineffective in enhancing covert spatial attention.

Another potential explanation for the absence of differences in alpha lateralisation between sham and active protocols is that external factors, e.g. screen dynamics, may have partially driven the observed alpha modulation. Feedback was delivered via a stimulus in the left visual field that dynamically changed colours, based on participants’ AAI, while the right stimulus remained black. Although this asymmetry was intended to minimise visual distractions, it may have introduced unintended noise into the EEG signal. Flashing or changing visual stimuli capture exogenous attention and elicit lateralised posterior alpha shifts that may have interfered with the ability to further modulate alpha power within the left and right hemispheres via neurofeedback (Casteau & Smith, 2020; Keefe & Störmer, 2021), while a dynamically changing stimulus might elicit evoked potentials causing sustained alpha lateralisation (Landry et al., 2024; Nathan & Hanley, 1975).

Finally, although behavioural effects have been reported from a similar neurofeedback protocol (Schneider et al., 2020), we found no differences in reaction times between active and sham neurofeedback sessions. The absence of both behavioural and neurophysiological effects further suggests that the neurofeedback task employed here was ineffective in modulating covert spatial attention.

### 4.2 tACS Does Not Shift Lateralised Alpha Power

Consistent with the behavioural results, we found no effect of session protocol or task timing on AAI values between the alpha and gamma tACS protocols. Although we hypothesised that tACS would induce a favourable brain state that would enhance neurofeedback performance, no lasting effects of alpha tACS were observed during the neurofeedback task nor post-stimulation Posner task. As mentioned above, aftereffects have been reported to persist beyond the time frame of our post-intervention Posner task (approximately 30–55 minutes after stimulation) (Orendáčová & Kvašňák, 2021). However, these durations are inconsistent and can be influenced by multiple factors, including electrode montage, stimulation frequency, stimulation duration, and pre-stimulation brain state (Veniero et al., 2015). While we found no effect of cue on the AAI signal in the Posner task, the general trend of results aligns with our initial expectations. Specifically, lower AAI values for right cues relative to left and centre cues indicate relatively higher alpha power in the ipsilateral hemisphere when attending to right cues, while similar power was observed across hemispheres for left and centre cues. This is in line with similar lateralisation of alpha power in covert spatial attention tasks (Kornrumpf et al., 2017; Peylo et al., 2021; Thut et al., 2006; Worden et al., 2000) and, coupled with the observed validity effect and topographies, further indicates that the participants were indeed performing the Posner task as instructed.

We also identified no effect of tACS on neurofeedback performance, as no differences in alpha power were observed between the alpha and gamma stimulation protocols immediately following tACS. Although growing evidence supports direct cortical entrainment as a mechanism of tACS, the exact underlying process remains debated (Asamoah et al., 2019; Elyamany et al., 2021; Modolo et al., 2024; Tavakoli & Yun, 2017). Furthermore, the optimal parameters for achieving entrainment, particularly for individual participants and specific experimental designs, are currently not well understood. In the absence of effective entrainment of alpha oscillations, we would not expect to have induced any differences in spectral power or reaction times following stimulation. Another possible explanation for the lack of tACS effects may be that the alpha lateralisation during the neurofeedback tasks was already at ceiling, and thus any entrainment achieved by tACS was negligible. However, similar tACS protocols were used previously to induce changes in reaction times on similar covert spatial attention tasks, which demonstrate an increase in lateralised alpha power (Kasten et al., 2020; Schuhmann et al., 2019). This suggests that the null results observed here may involve additional contributing factors beyond ceiling effects.

### 4.3 No Impact of tACS Stimulation on Reaction Times

We hypothesised that tACS would enhance the ability to modulate left-directed covert spatial attention during neurofeedback by entraining individual alpha frequency in the ipsilateral (left) posterior hemisphere and, through interhemispheric competition, a reduction of alpha power in the right hemisphere (Gallotto et al., 2020; Kinsbourne, 1977). Given the established link between covert spatial attention and lateralised posterior alpha power (Kornrumpf et al., 2017; Peylo et al., 2021; Rihs et al., 2007; Sauseng et al., 2005; Thut et al., 2006), we anticipated that alpha tACS would reduce reaction times to valid left targets and increase reaction times to valid right targets in the Posner task compared to gamma tACS. However, we observed no differences in lateralised reaction times to valid cues or in overall reaction times to valid and invalid cues across the three tACS protocols. The lack of significant findings suggests that tACS did not modulate neurofeedback performance or the associated behavioural outcomes.

Previous studies have reported changes in reaction times during endogenous attention tasks following similar tACS protocols, even in the absence of neurofeedback (Kasten et al., 2020; Kemmerer et al., 2022; Schuhmann et al., 2019). Although these studies differed as they applied tACS during the tasks and used various electrode montages, there is evidence that tACS aftereffects, such as entrained power differences, can persist for tens of minutes after stimulation (Orendáčová & Kvašňák, 2021; Veniero et al., 2015). Furthermore, singular, pad electrodes, like those used in our study, have been shown to induce measurable oscillatory and behavioural effects (Gundlach et al., 2017; Hopfinger et al., 2017; Kasten et al., 2016; Vossen et al., 2015). Despite these findings, null results have also been reported with similar protocols (Brignani et al., 2013; Coldea et al., 2021; Silas et al., 2023).

Several factors may influence tACS efficacy, including electrode montage size and position, stimulation intensity, duration, frequency, and the number of sessions (De Koninck et al., 2023; Orendáčová & Kvašňák, 2021; Tavakoli & Yun, 2017). In addition, the endogenous brain state prior to stimulation is likely to influence its efficacy (Bradley et al., 2022; Kasten & Herrmann, 2022; Ruhnau et al., 2016). Specifically, tACS may only be effective when endogenous oscillations are engaged but not already at ceiling levels (Ruhnau et al., 2016), and may be limited to specific brain states that are continuously fluctuating during stimulation (Kasten & Herrmann, 2022). While the occipito-parietal alpha power in our study was unlikely to be at ceiling and should, in principle, have been susceptible to tACS entrainment, the complex and dynamic interaction between ongoing brain states and tACS efficacy remains uncertain. To date, no systematic investigation has fully categorised the multiple interactions between non-invasive brain stimulation parameters and ongoing brain states, making it difficult to determine which factors may facilitate or hinder tACS efficacy. This lack of clarity may partially account for the null findings observed in this study and raises broader questions about the suitability of tACS as a tool for neural modulation and entrainment.

Despite the lack of effect from tACS stimulation, the observed validity effect confirms that participants successfully engaged with the Posner task. Specifically, participants consistently responded faster to targets appearing on the cued side, indicating successful shifts of attention in response to the directional cues (Posner, 1980). This pattern is supported by the posterior lateralisation of alpha band topography, with increased ipsilateral alpha power and decreased contralateral alpha power relative to the cue direction (Kornrumpf et al., 2017; Peylo et al., 2021; Thut et al., 2006; Worden et al., 2000).

### 4.4 Questionnaire results

Despite the lack of behavioural and neurophysiological effects, it remains possible that participants were able to distinguish between the active neurofeedback and sham sessions. If so, this may indicate that our measures were not sensitive enough to detect subtle effects, or could reflect factors such as de-motivation, or nocebo effects, arising from a belief that they were not controlling their brain activity (Geers et al., 2005; Schönenberg et al., 2021). To explore this possibility, we analysed the end-of-session questionnaires. Firstly, participants were unable to identify the sham neurofeedback session at a rate higher than chance, suggesting that they were generally unaware of the condition that was administered. Although, we did not probe whether they could identify the alpha from the gamma tACS stimulation, we reasoned that this information would not have influenced behaviour since most participants would not have had prior knowledge of the expected effects of different stimulation frequencies. It is also possible that the positive feelings of subjective success may be linked to a general placebo effect associated with participating in a neurofeedback and tACS experiment, regardless of condition (Geers et al., 2005). To further explore this, we compared the perceived success and motivation of those participants who could identify the sham neurofeedback compared to those who could not. We also investigated motivation and perceived success between session protocols, but found no significant differences across these groups, further supporting the conclusion that participants were unaware of their assigned condition.

### 4.5 Limitations

There are several limitations to this study. Firstly, although rectangular pad electrodes have been shown to elicit tACS effects, ring electrodes may provide a more focused delivery of the electric field towards the cortex (Herrmann et al., 2013). However, since we aimed to minimise tACS artefacts in the EEG signal, particularly around the neurofeedback electrode sites, we chose to proceed with smaller, pad electrodes because they covered a smaller and moderately focal area of the scalp. Alternative electrode montages that provide greater focality than the conventional pad-based Cz–O1 configuration used here, such as HD-tACS setups employing multiple smaller electrodes, may improve electric field targeting while reducing EEG artefacts (Wu et al., 2021). We also applied the tACS offline, before the neurofeedback training, due to artefact contamination in the EEG feedback signal when tACS was delivered online, during neurofeedback. If any tACS priming effects were present but short-lived, this would not have had any facilitatory effect during the subsequent neurofeedback period. Future developments in online artefact removal may make combined tACS and neurofeedback more feasible and effective.

Secondly, many neurofeedback paradigms involve multiple training sessions to enable participants to become proficient with the task and to capture more subtle, sustained learning effects. Although we employed a within-subject design with three sessions per participant, each session differed in protocol, and no constraints were placed on the time interval between sessions. This variability may have enabled learning in some participants and limited learning in others. Further, while it could be argued that a single session is insufficient for participants to achieve proficiency in the neurofeedback task, the CVSA paradigm was designed to facilitate learning without extensive training blocks or reliance on self-generated strategies, factors often implicated in neurofeedback control variability (Chen et al., 2022; Wang et al., 2023). Still, it may be unrealistic to expect improvements in covert spatial attention even within three sessions and more sustained, repeated training may be needed, e.g. 20-30 sessions (Zoefel et al., 2011), particularly if similar protocols were to be tested in clinical groups such as patients with hemispatial neglect.

### 4.6 Conclusion

Our findings indicate that individual alpha frequency tACS did not affect neurofeedback performance, as measured by changes in reaction times in the Posner task or alterations in alpha-band lateralisation following stimulation. However, a robust validity effect confirmed that participants were orienting covert attention as instructed, supporting the interpretation that the accompanying posterior alpha lateralisation reflected attentional modulation associated with the task. Similarly, active neurofeedback did not induce measurable changes in alpha lateralisation or behavioural performance compared to sham neurofeedback, although alpha activity was modulated in a manner suggestive of leftward covert spatial attention engagement in all 3 experimental conditions. Our findings suggest that posterior alpha tACS, applied prior to neurofeedback, does not enhance neurofeedback performance in the context of our covert spatial attention paradigm. Nevertheless, tACS may influence neurofeedback outcomes in other paradigms or modalities, highlighting a need for rigorous studies investigating the effects of different tACS parameters on behavioural and neurophysiological outcomes.

## 5 Statements and Declarations

## 5.1 Data and Code Availability

Raw data in BIDS format in addition to scripts for analysis and visualisation are available on the Open Science Framework (OSF) (Turner et al., 2022).

## 5.2 Author Contributions

C.T.: Conceptualisation, Methodology, Software, Formal analysis, Investigation, Data Curation, Writing - Original Draft, Writing - Review & Editing, Visualisation, Project administration, Funding acquisition

L.J. Investigation, Writing - Review & Editing,

M.B. Investigation, Writing - Review & Editing,

A.V. Supervision, Conceptualisation, Writing - Review & Editing, Funding acquisition

G. L. Supervision, Conceptualisation, Methodology, Resources, Writing - Review & Editing, Funding acquisition

A.F. Supervision, Conceptualisation, Software, Resources, Writing - Review & Editing, Funding acquisition

## 5.3 Funding

C.T. was supported by an Engineering and Physical Sciences Research Council (EPSRC) studentship (EP/T517896/1)

A.V. Engineering and Physical Sciences Research Council (EPSRC) studentship (EP/T517896/1)

G.L. was supported by the Wellcome Trust [209209/Z/17/Z].

A.F. was supported by a grant from the Biotechnology and Biology Research Council (BBSRC, grant number: BB/S006605/1) and the Bial Foundation (Bial Foundation Grants Programme; Grant id: A-29315, number: 203/2020, grant edition: G-15516).

